# Monkey hear, monkey do what? An application of Automated Behavioural Response systems for hypothesis testing in the world’s smallest monkey

**DOI:** 10.64898/2026.05.28.728389

**Authors:** Larissa Barker, Sarah Papworth

## Abstract

Observer effects are a frequent problem in animal behaviour studies, particularly when assessing responses to human disturbance. Automated Behavioural Response (ABR) systems, which combine camera traps with automated sound playbacks, offer a solution but have been primarily used on large terrestrial mammals. Here, we demonstrate their use in a small (∼110g) arboreal primate, the eastern pygmy marmoset (*Cebuella niveiventris*). We conducted two playback experiments to test the risk-disturbance and distracted prey hypotheses. The marmosets exhibited strong anti-predator responses to avian predator calls, including increased fleeing and vocalisations. Human speech elicited similar but weaker responses, indicating that pygmy marmosets do not perceive raptors and humans as equivalent threats. Embedding predator calls into anthropogenic noise reduced vocal responses, suggesting that anthropogenic noise interferes with responses to predation cues. Across five weeks, we generated 128 successful experimental trials, demonstrating that ABRs can rapidly produce sample sizes sufficient for hypothesis testing in the field.

## Introduction

Audio playback experiments allow a researcher to create a naturally rare event and are a vital instrument for understanding animal behaviour (Tyack, 2009). Historically, these experiments have been conducted by placing a speaker out of view of the animal, with a researcher recording the focal individual’s reaction to the audio stimuli (Fischer et al., 2013). However, researchers being present while recording data can cause observer effects, where the study organism shifts their behaviour due observer presence (McDougall, 2012). These observer effects have been found to bias behavioural studies (Jack et al., 2008). One promising new technological development is the Automated Behavioural Response (ABR) system (Suraci et al. 2017). The system combines a camera trap with a speaker, which is programmed to play an audio recording when the camera trap is triggered (Suraci et al., 2017). This non-invasive approach records animal responses to audio playbacks, so can be used to study questions such as predator-prey interactions and anti-predator responses in wild animals (Suraci et al., 2017). The ABR is a powerful tool which allows researchers to capture the responses of focal species to a variety of audio cues. The ABR system has primarily been used to describe species interactions (Suraci et al., 2017; Epperly et al., 2021, Rigoudy et al., 2022), including understanding wildlife responses to anthropogenic noise and human speech (Smith et al., 2017; Mugerwa, 2018; Zeller et al., 2024). Recent deployments of ABRs have shown their potential use for reducing human wildlife conflict by reducing wildlife visitation to targeted areas (Bhardwaj et al., 2022), as a tool for nonlethal wildlife management (Yiu et al., 2024) and as a mechanism to reduce crop damage by ungulates (Widén et al., 2022).

ABRs have also been suggested to have the unique ability of generating sufficient sample sizes to statistically evaluate ecological and behavioural hypotheses that were previously untestable due to the high field time costs of former ‘in-person’ playback methods (Suraci et al., 2017; Rigoudy et al., 2022; Zeller et al., 2024). However, ABRs have not yet been applied to hypothesis testing, and have focused on large (>40kg), terrestrial mammals in open habitats (Fletcher et al., 2023; Kasper et al., 2025). The large sample sizes potentially generated by ABRs may make them particularly suitable for studying animal behaviour in tropical rainforest environments, as most of the wildlife found in these habitats occur at low densities and are naturally cryptic (Linkie et al., 2008).

Here, we demonstrate the targeted use of ABRs for conducting hypothesis-testing playback experiments on a small (∼110g) arboreal species in a tropical rainforest. This study is the first direct test of described behavioural hypotheses with the ABR system, the first use in a solely arboreal environment, as well as the first application of the system to an animal smaller than 500g (the smallest animal previously studied with ABRs is the fox squirrel (*Sciurus niger*), Potash et al., 2023). Specifically, we designed two playback experiments to test specific hypotheses about eastern pygmy marmoset (*Cebuella niveventris*) responses to humans and predators. The first experiment tested the risk-disturbance hypothesis, which postulates that animals perceive human disturbance similarly to predation risk (Walter, 1969; Frid and Dill, 2002). A lack of difference in responses to playbacks of human sounds and predators, and increased responsiveness compared to control playbacks, would support this hypothesis. When compared to control playbacks, expected pygmy marmoset responses to predator calls are fleeing, increased vigilance and vocalisations (Snowdon and de la Torre, 2002; Ferrari and Ferrari, 1990). Under the risk-disturbance hypothesis, we would therefore expect the same responses to birds of prey as to our anthropogenic playbacks of human speech and motor boats, and no change in behaviour in response to control playbacks of cicadas and macaws. The second experiment explored the distracted prey hypothesis, which postulates that animals are distracted by any stimulus they can perceive, and this distraction can cause them to become more susceptible to predation (Chan et al., 2010). We played audio recordings of motor boats and human speech that were spliced with the calls of birds of prey and control audios. The distracted prey hypothesis would be supported if pygmy marmosets show reduced responsiveness to predator playbacks when spliced into playbacks of motor boats and human speech.

## Results

We deployed ABR systems focused on the feeding trees of nine pygmy marmoset groups during the 5-week study. Four of the groups generated just seven successful trials between them, even though 183 videos were recorded across these groups (Table 1). For the other five groups however, we recorded 1,268 videos and 128 successful experimental trials in a 5-week study, demonstrating that the ABR system can be used to generate data for hypothesis testing. For these five groups we could evaluate their responses to the stimuli for both experiments, and below we summarize the results of the trials.

**Table 1:** A breakdown of the trials each group underwent in the first experiment.

| Included in the final analysis | Group ID | Total number of videos recorded | Total number of deployments | Total number of successful trials | Control with no playback | Control playback | Motor Boat playback | Human Speech playback | Avian predator call playback |
| --- | --- | --- | --- | --- | --- | --- | --- | --- | --- |
| No | CV3 | 105 | 1 | 0 | 0 | 0 | 0 | 0 | 0 |
|  | RC2 | 41 | 2 | 3 | 0 | 1 | 0 | 0 | 2 |
|  | RC7 | 23 | 2 | 2 | 0 | 0 | 1 | 1 | 0 |
|  | TL11 | 19 | 3 | 2 | 0 | 0 | 0 | 1 | 1 |
| Yes | CV2 | 104 | 1 | 10 | 14 | 2 | 2 | 2 | 4 |
|  | CV4 | 112 | 1 | 9 | 14 | 2 | 1 | 3 | 3 |
|  | CV7 | 51 | 2 | 20 | 14 | 5 | 4 | 5 | 6 |
|  | TL7 | 15 | 1 | 6 | 14 | 1 | 1 | 2 | 2 |
|  | TL14 | 26 | 1 | 9 | 14 | 2 | 2 | 2 | 3 |
|  | Total included |  |  |  | 70 | 12 | 10 | 14 | 18 |

### Experiment 1: The risk disturbance hypothesis

Fifty-four playback trials were generated for the first experiment, and we augmented this with 70 videos without playbacks which were used as a control (Table 1). Avian predator call playbacks had the highest percentage of trials where an individual vocalised, 72% (Figure SM1 in supplementary materials). Motorboat playbacks had the next highest percentage of trials with vocalisations at 50%. Overall, the number of vocalisations differed between conditions (Anova Type II_4_, X^2^=15.31, p=0.004, n=124), with 6 (95% CI: 2.5%=2.25, 97.5%=14.75) more vocalisations in videos with avian predator playbacks compared to videos with no playback (GLMM, SE= 2.76, z=3.65, p<0.001, distribution=negative binomial, n=124). No other playback conditions had a significant effect on the number of vocalisations when compared to videos with no playback.

The focal individual was significantly more likely to flee in playback trials with avian predator calls (β = 0.32, t = 4.33, permutation p<0.001, n=124) and human speech (β = 0.27, t = 3.33, permutation p = 0.002, n=124) (Figure 1) when compared to videos with no playback. There was no significant effect on fleeing likelihood for playbacks with motor boat (β = 0.09, t = 0.95, p = 0.27, n=124) or control sounds (β = –0.006, *t* = –0.07, *p* = 0.93, n=124). After fleeing occurred, the mean time for any pygmy marmoset to return to the recording area after an avian predator call playback was 74.80 minutes (n=6), 54.08 minutes (n=4) for human speech playbacks, 0.90 minutes (n=2) for no playback trials and 2.75 minutes (n=1) for motor boat playbacks (Figure 2, Supplementary materials Table SM1).

**Figure 1.**
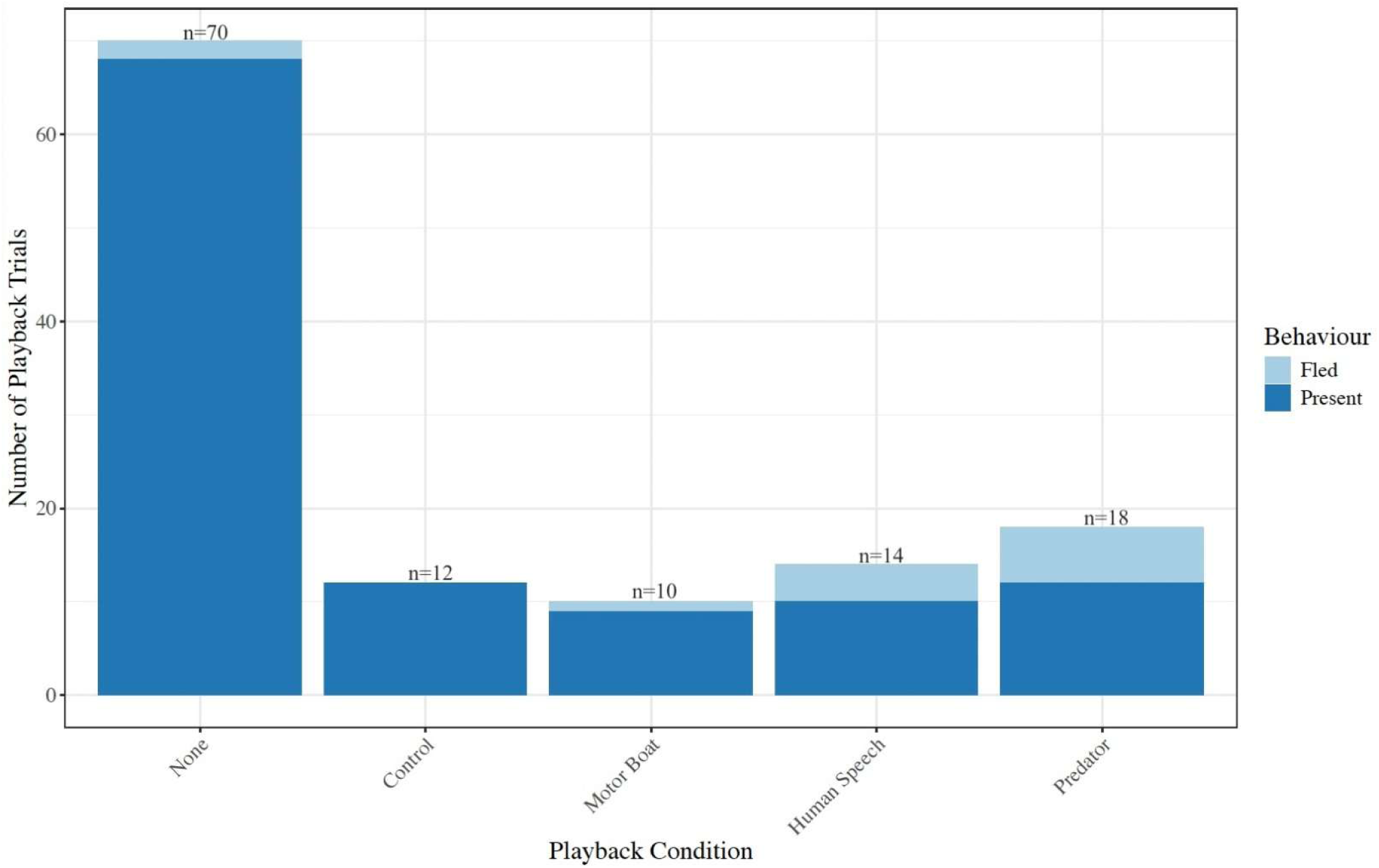
The number of trials where the focal individual fled during a playback condition (x axis) and number of trials (y axis) where they were present for the duration of the trial (includes trials where the focal individual did move out of the video frame but not because they were fleeing). The number of successful trials for each condition is shown above each bar.

**Figure 2:**
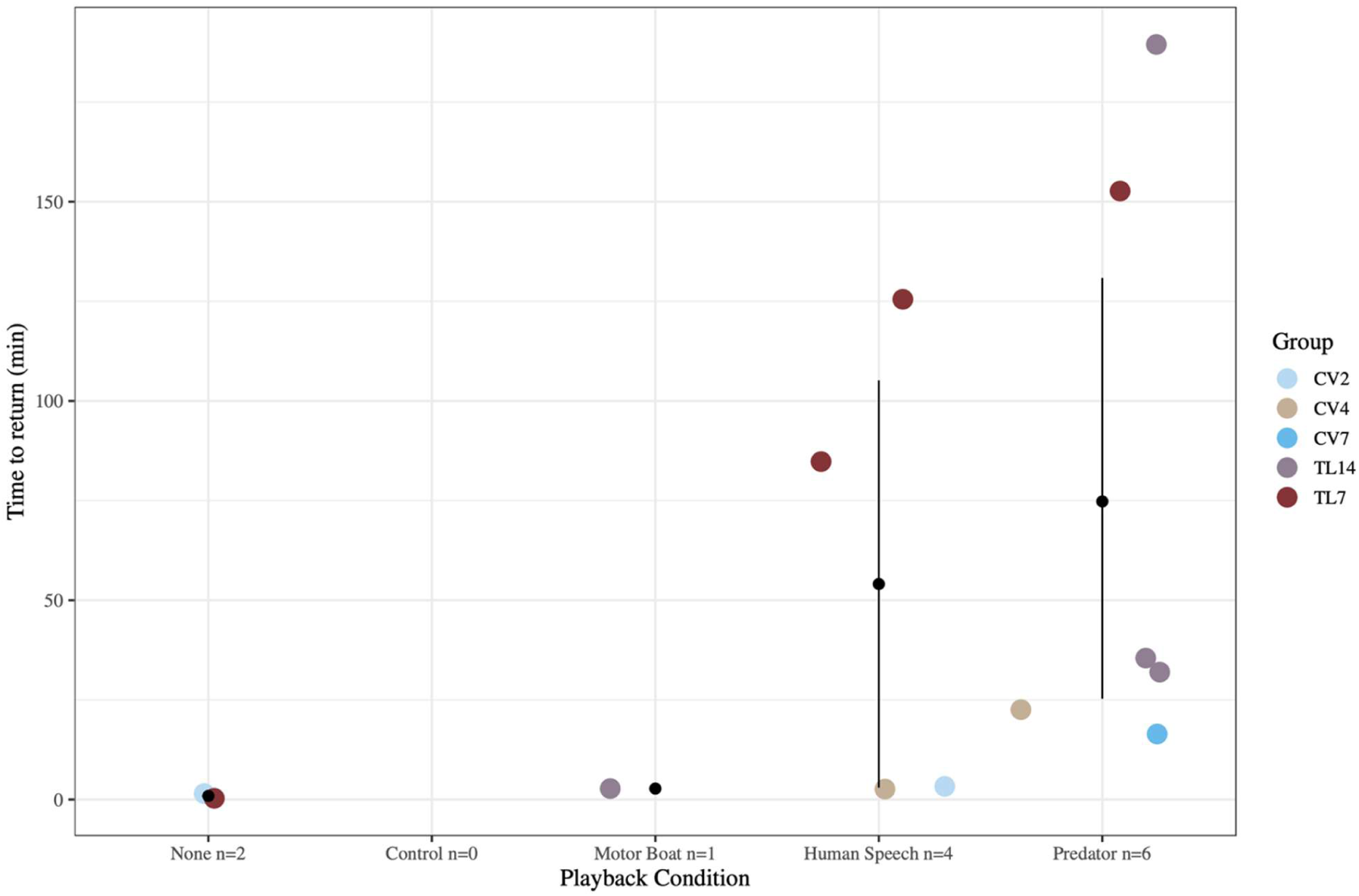
After a fleeing event, the time in minutes until a pygmy marmoset triggered the camera trap again. The black dot indicates the mean time to return, and confidence limits are shown for conditions where there are more than two fleeing events.

Playback condition did not have a significant effect on the duration of eating (Anova Type II_4_, X^2^=8.80, p=0.07, n=124), vigilance (Anova Type II_4_, X^2^=1.77, p=0.78, n=124) or looking at the camera trap (Anova Type II_4_, X^2^=3.52, p=0.48, n=124). In all three models there was a significant positive relationship between time in frame (TIF) and the three behaviours, suggesting that the behaviours had longer durations when individuals were visible for longer (Anova Type II_1_, X^2^=130.95, p<0.001, n=124; Anova Type II_1_, X^2^=257.43, p<0.001, n=124; Anova Type II_1_, X^2^=23.50, p<0.001, n=124; respectively).

### Experiment 2: The distracted prey hypothesis

A total of 74 videos were generated for the second experiment (Table 2). Overall, the number of vocalisations did not differ between the human speech and motor boat playback conditions (Anova Type III_1_, X^2^=3.07, p=0.08, n=74; supplementary materials Figure SM1). There was also no effect of which audio was spliced in (control or avian predator call) (Anova Type III_1_, X^2^=0.73, p=0.39, n=74) nor an interaction between the two terms (Anova Type III_1_, X^2^=1.82, p=0.18, n=74).

**Table 2:** The number of trials for each condition and group in the second experiment.

| Group ID | Total number of videos recorded | Total number of deployments | Total number of successful trials | Motor Boat + Controls | Motor Boat + Predator | Human Speech + Controls | Human Speech + Predator Calls |
| --- | --- | --- | --- | --- | --- | --- | --- |
| CV2 | 93 | 2 | 17 | 3 | 3 | 2 | 9 |
| CV4 | 201 | 3 | 12 | 1 | 4 | 3 | 4 |
| CV7 | 16 | 1 | 10 | 3 | 2 | 1 | 4 |
| TL7 | 196 | 3 | 14 | 4 | 4 | 0 | 6 |
| TL14 | 266 | 2 | 21 | 3 | 6 | 5 | 7 |
| Total included |  |  |  | 14 | 19 | 11 | 30 |

To examine the predictions of the distracted prey hypothesis, we compared the number of vocalisations in response to the spliced predator calls playbacks to the number of vocalisations in response to the predator playback condition in the first experiment. We found that audio condition did have a significant effect on the number of vocalisations emitted (Anova Type II_2_, X^2^=20.28, p<0.001, n=67). There were 2 (95% CI: 2.5%=0.60, 97.5%=7.98; Supplementary materials Figure SM1) more vocalisations in the predator only playback experiments compared to the playbacks when predator calls were spliced into either anthropogenic noise (GLMM, distribution=negative binomial, n=67, Human speech spliced with predator; estimate= 0.17, 95% CI: 2.5%=0.07, 97.5%=0.40, z=-3.95, p<0.001, n=67; Motor boat spliced with predator, estimate= 0.15, 95% CI: 2.5%=0.05, 97.5%=0.42, z=-3.62, p<0.001).

No fleeing was observed after the motor boat spliced with control sound playback, whereas fleeing was observed in response to all other conditions (Figure 3). Focal individuals were significantly more likely to flee during human speech audio playbacks compared to motor boat playbacks (β = 0.27, *t* = 1.76, permutation p=0.001, n = 74). The presence of a predator call spliced condition marginally increased fleeing likelihood overall (β = 0.15, *t* = 1.11, permutation p = 0.06, n = 74) (Figure 4). There was also a significant interaction between anthropogenic audio type and splice condition (β = –0.19, *t* = –0.99, permutation p = 0.008, n=74) which suggests these effects are not additive (Figure 3). After fleeing occurred, the mean time to return after a human speech audio spliced with a predator audio was 34.68 minutes (n=7), after a motor boat audio spliced with a predator audio was 30.18 minutes (n=3) and after a human speech audio spliced with a control audio 15.34 minutes (n=3) (Figure 4, Supplementary materials Table SM1).

**Figure 3.**
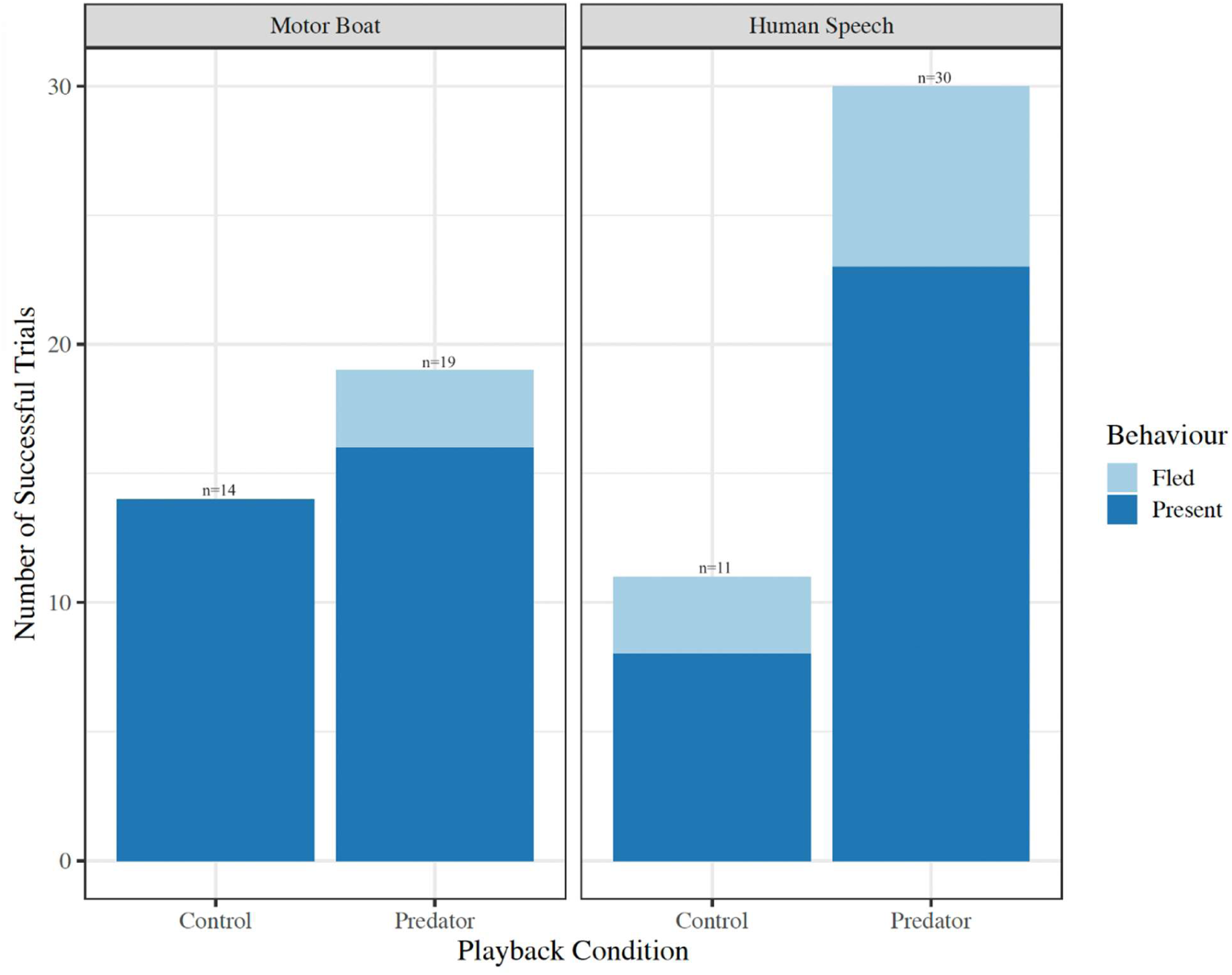
The total number of playbacks where the focal individual fled and playbacks where they were present during the duration of the trial (includes playbacks where the focal individual did move out of the video frame but not because they were fleeing). Each main anthropogenic noise condition (human speech and motor boat) is shown separately, with trials split into those where these were spliced with a control sound or a predator call. The number of successful playbacks is shown above each bar.

**Figure 4:**
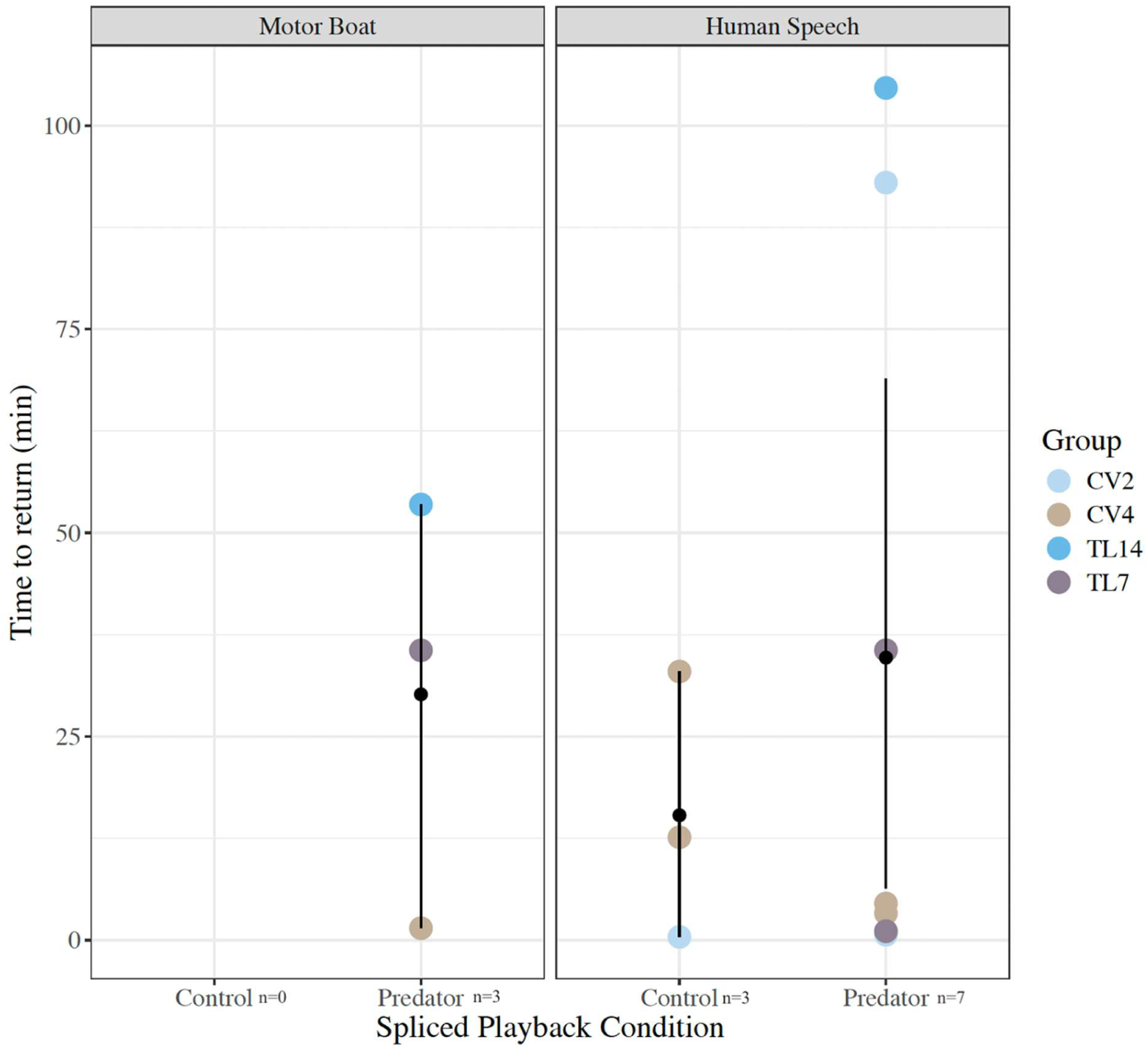
After a fleeing event, the time in minutes until another marmoset triggered the camera trap again. The black dot indicates the mean time to return with confidence limits for that audio condition.

Eating duration was not correlated with the either anthropogenic noise in the playback (Anova Type III_1_, X^2^=0.19, p=0.67, n=74), the audio that was spliced in (Anova Type III_1_, X^2^=1.33 p=0.25) or the interaction between them (Anova Type III_1_, X^2^=0.02, p=0.88, n=74). Eating duration only correlated with TIF, showing that longer eating durations were observed when the individual was visible for longer (Anova Type III_1_, X^2^=43.52, p<0.001, n=74). This was the same for duration of vigilance behaviour, with only TIF having a significant effect (Anova Type III_1_, X^2^=81.54, p<0.001, n=74).

## Discussion

We first discuss the results of the two experiments, and then the application of ABR technology demonstrated by this study.

### Understanding pygmy marmoset responses to humans and predators

We find partial support for both hypotheses, as responses to human sounds were similar to responses to predators, and the response to predators was less intense when the sound was spliced into anthropogenic sound.

Our results from the first experiment on the risk disturbance hypothesis partially align with our predictions: predator calls resulted in the most fleeing events and contained the most vocalisations. Human speech playbacks had the second highest percentage of fleeing events and second longest return times, which aligns with one previous study which found pygmy marmosets moved out of sight when played loud recordings of human speech (Sheehan and Papworth, 2019). The results of this study differ from Hawkins and Papworth (2022), who found no behavioural changes in response to playbacks of human speech, and therefore concluded there was little evidence to support the risk-disturbance hypothesis in pygmy marmosets. In contrast, the fleeing behaviour in this study provides partial support for the risk-disturbance hypothesis, which predicts similar responses to humans as to predators, as both have significantly higher fleeing rates. However, we did not find identical responses, as frequency of vocalisations was higher after predator playbacks than in the no playback videos, but not after human speech playbacks. While pygmy marmosets might perceive both raptors and humans as predators, they may have different anti-predator responses to these threats. Little is known about pygmy marmoset predation and anti-predator behaviour so we cannot say for certain if pygmy marmosets, like other species of primates, have predator specific behaviour (Fichtel et al., 2005). We did not detect any behavioural changes in the duration of eating or vigilance between playback conditions.

The differences found in our study and those in Sheehan and Papworth (2019) and Hawkins and Papworth (2022) could be because both those studies had a human observer present, influencing the responses by pygmy marmosets. The second experiment showed some support for the distracted prey hypothesis, suggesting that humans may affect responses to predator. We found predator calls resulted in fewer fleeing events when embedded within a playback of human speech. Surprisingly though, fleeing rates were similar between predator-spliced and control-spliced human speech conditions, even though both human speech and predator calls, when presented alone in the first experiment, elicited high levels of fleeing. Individuals still fled more often during human speech playbacks compared to motor boat playbacks, regardless of what was spliced in. This suggests greater responsiveness to human speech than motor boat noise. Pygmy marmosets gave around 2 more vocalisations in the predator call condition from experiment 1 when compared to the spliced playbacks in experiment two, indicating some impact of anthropogenic noise on vocalisation. However, as this masking is specific to one type of anthropogenic noise (human speech) our results did not fully support the risk-disturbance hypothesis.

All five groups included in this study had relatively high levels of exposure to anthropogenic noise (Barker, 2022). Anti-predator behaviours are an important survival tool that have population level consequences (West et al., 2017), so testing the responses of less exposed pygmy marmoset groups is necessary before generalizations can be made to the broader population. Our results show that even pygmy marmosets with higher human exposure were more likely to flee from feeding holes and delay their return when they heard a playback of a avian predator call or human speech. If these disturbances occur frequently, it could have a substantial impact on the feeding behaviour of these primates. As these results suggest human speech does have an impact on pygmy marmoset behaviour, we suggest that tour operators reduce stress for the animals by informing their guests to be silent while approaching and viewing the monkeys.

### Use of ABR systems for small arboreal species and hypothesis testing

We establish that ABR systems can be applied in arboreal environments and for far smaller species than previous demonstrated, expanding the future applications of this technology. Our study also demonstrates the usefulness of ABRs as a tool for field hypothesis testing. This study was constrained by Covid-19 travel restrictions, limiting the number of groups we were able to include in our analyses. Nevertheless, in five weeks we were able to collect 198 videos which could be analysed to test two hypotheses, including 128 experimental playback trials. This is more than three times the data points collected by two previous playback studies on the same species with longer field periods (Hawkins and Papworth, 2022; Sheehan and Papworth, 2019). Therefore, we provide crucial evidence to support the statements made by Palmer et al. (2022) and Smith et al. (2020) that the ABR can generate sufficient sample sizes to evaluate ecological and even broader behavioural hypotheses. While the pygmy marmoset is a particularly amenable species for ABR techniques due to their regular return to specific feeding trees to extract sap and gum from pre-existing holes, the ABR system has previously been used successfully in a wide variety of mammalian species (Smith et al., 2017; Mugerwa, 2018; Widén et al., 2022).

This study also provided invaluable field insights for future deployments in similar systems. Our first recommendation is the use of brief pilots for specific study designs, study species and sites to allow an estimation of the number of deployments and field days to generate sufficient sample sizes. Although ABRs can generate large sample sizes, study designs with too many experimental conditions may fail to generate a large enough sample per condition if the target species triggers the camera too infrequently. A second recommendation is to erect the equipment without batteries (so the ABR is not triggered) for 2-3 days before experiments start, to ensure animals are used to the presence of the system in their environment. Although camera traps are more passive and eliminate the need for a human observer, some animals still find their presence intrusive (Meek et al., 2016). Two out of the four groups that were not included in the final analysis of both experiments were apprehensive of the camera and especially the speaker, which is why they generated so few successful playbacks.

There are limitations to the ABR methodology which should be considered before committing to their use in future studies. The automated function means experiments can be conducted without human presence but is logistically harder to make sure that each group undergoes the same number of playbacks. This can lead to over- or under- representation of some playback conditions for certain groups. Specific individuals within a group may also be overrepresented in the videos, as individuals are not identified. Without a human present to identify or track individuals, we cannot be sure that the same individual is not continuously returning. This was not a major limitation for these experiments as pygmy marmoset groups do not share feeding holes so we considered the group as our unit of analysis. However, these limitations may mean that the ABR will not be the most appropriate methodology for all behavioural studies. One technical challenge in this study was the advanced electrical engineering knowledge required to create an ABR system, but the development of the ‘BoomBox’ by Palmer et al. (2022) - an open-source Arduino-compatible board which one can attach to any commercially available camera trap to form an ABR system - makes sourcing an ABR more cost and time efficient.

While this study shows the feasibility of using ABRs to test animal behavioural hypotheses in the field, analysis is time-consuming. Across the two experiments, 1,268 videos were produced that then needed to be reviewed to see if there was a successful playback. Of these 1,268 videos, 198 were then analysed in BORIS. Camera trap technology expands sample sizes but also generates comprehensive audio-visual datasets that are often so large and data-rich that a timely categorisation and analysis by human researchers is no longer possible (Bain et al., 2021). A potential future avenue to bypass this limitation would be to integrate machine learning analytical methods to review and classify videos. Machine learning, artificial intelligence and deep learning have all seen a recent spike in popularity (Pichler and Hartig, 2023) and are revolutionising a wide scope of scientific studies (Jordan and Mitchell, 2015). The integration of these automated data processing algorithms is at the forefront of ethology and ecological studies (Saoud et al., 2024). Further developing these machine and deep learning algorithms would allow for a widespread application of the ABR systems to test a variety of behavioural hypotheses in this and other systems.

Our application of the ABR system with a small arboreal primate demonstrates how this methodology can be applied in future behavioural ecology studies. Using the recommendations discussed above, the ABR has the potential to transform how playback experiments are conducted. The ABR system allows researchers to test behavioural hypotheses, including those dealing with the impacts of human disturbance, without the bias created by human observers. Eliminating this bias allows researchers to fully understand the impacts of human presence. This can aid studies on human impacts, leading to practical mitigation efforts, and inform broader behavioural studies by generating more ecologically relevant observations of animal behaviour.

## Methods

### Study Site

This research was conducted in the Área de Conservación Regional Comunal Tamshiyacu Tahuayo (ACRCTT) a communal reserve located in north-eastern Peru. Established in 1991 by the local community, researchers and conservationists, the reserve was upgraded to a state reserve in 2009 and increased to 420,080 ha (Penn, 2009). Tourists can visit the reserve by staying at Amazonia Expedition’s main accommodation, just out of the reserve limits and close to the community of El Chino (Figure 5). The only manmade structure inside the reserve is a research centre, also run by Amazonia Exhibitions. Their peak tourist season is July-August, during the dry low-river season. The main source of anthropogenic noise in the area comes from motor boats (Barker, 2022), the primary transport in the area. There are significant differences in the amount and variety of anthropogenic noise inside versus outside the reserve, with the majority of the anthropogenic noise coming from El Chino and the main lodge (Barker, 2022).

**Figure 5.**
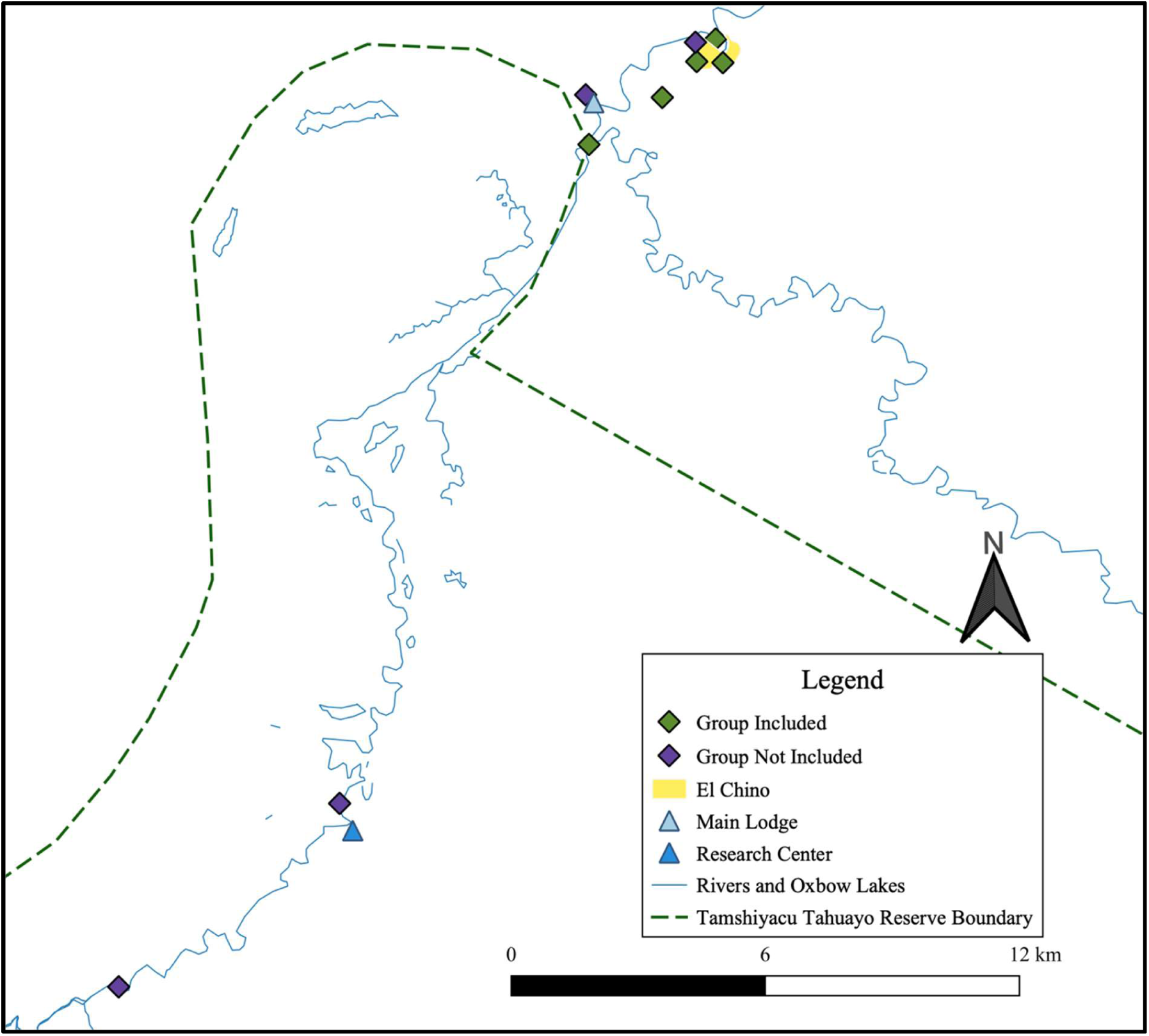
A map of the locations of the marmoset groups where the ABRs were placed in September to October 2021. Green diamonds denote groups that underwent successful trials and were included in the final analysis. Groups that did not complete enough playbacks to be included in the final analysis are shown as purple diamonds.

### Study Species

The eastern pygmy marmoset, *Cebuella niveiventris*, is a 110-120g Neotropical primate species, a habitat specialist found in the forests along river-edges in the Amazon rainforests of Bolivia, Brazil, Ecuador and Peru (de la Torre et al., 2021). Pygmy marmosets are diurnal and active from just after sunrise to just after sunset, ∼05:30-18:30 (Ramirez *et al*., 1977). Group size ranges from 2-9 individuals (de la Torre et al., 2000). Home ranges for a group are between 0.1 to 0.5 ha, and feature 1-6 central feeding trees where they create holes to extract sap and gum (Soini, 1988). Their small home ranges and repeated visits to a small number of trees make it relatively easy to predict their daily movements (de la Torre et al., 2000), making them popular with tour operators. As a specialist species, they are also vulnerable to changes in habitat and human activity (de la Torre et al., 2000). They are also sensitive to capture and pet trade trafficking, which causes behavioural change and has been found to decrease group size and reproductive rates (de la Torre et al., 2000).

We focused on five groups, as a further four groups had to be excluded due to the low encounter rates on the camera traps. Anthropogenic noise, including human speech and motor boats were monitored for three of these five groups, showing they were exposed to an average 20 hours and 29 minutes of anthropogenic noise over 48 hours (Barker, 2022). We did not conduct follows to determine full group composition for these groups as this study is a part of a larger research project on the impacts of human exposure on pygmy marmosets, so we did not want to bias future studies by habituating groups. Groups were chosen if they had feeding holes in positions that could be captured on the ABR, and none of these groups had previously taken part in an ABR playback experiment. Two of the five groups were included in a playback experiment in 2017 (Sheehan and Papworth, 2019) and three groups in 2019 (Hawkins and Papworth, 2022).

### ABR set up

The 4 ABR units used in this study were constructed using the specifications outlined in Suraci et al. (2017) and altered to fit the parameters of this study. The ABR system includes a camera trap, mp3 player, and a speaker. Over 5-weeks in September – October 2021 (the dry season), an ABR was placed focusing on active feeding holes of 9 pygmy marmoset groups, which were identified during preliminary site visits. The ABRs were attached to an adjacent tree between 1-2 meters away from the feeding tree. The speaker was positioned 0-1.5 meters from the ABR and concealed and camouflaged with leaves or other greenery to help disguise its presence from the marmosets (Supplementary materials Figure SM2).

10 seconds after the camera trap detected movement and started recording a 2-minute video, a 30 or 60 second audio recording stored on an MP3 player was triggered (see stimuli details below) and started playing on the attached speaker. After a stimuli was played, there was a holdoff period of 12 minutes, where further triggers of the camera trap would not prompt additional playbacks. After the 12-minute holdoff, the next camera trap trigger would prompt the MP3 player to play the next track. As pygmy marmosets are diurnal, the ABRs were set up in the morning before 07:30 and set to hibernate after being active for 12 hours so the speaker would not continue playing while the marmosets were inactive.

After two days the equipment was removed, and the number of successful playbacks was reviewed. Successful playbacks were those where a video was recorded with an individual marmoset visible in frame before and at the start of the playback. If there were not enough successful playbacks for one or more conditions, then after a two-day break the ABR was redeployed at the same location with only the missing conditions. As individuals were not habituated, we do not know which individuals within the group experienced the trials, and thus we considered the group, rather than the individual, our unit of analysis. Of the 9 groups, 5 had successful playback trials for all conditions in the first experiment and were included in the final analysis for both experiments (Figure 5).

### Experimental procedure

The first experiment measured reactions to single stimuli and tested the risk-disturbance hypothesis. The experiment included the analysis of videos where no sound was played and those where 30 seconds of continuous anthropogenic noise (human speech or motor boats), predator calls and control sounds (see below) were played to pygmy marmosets. The second playback experiment tested the distracted prey hypothesis and extended the length of the playback to 60 seconds. Anthropogenic noise, either human speech or motorboat, was played for one minute and after 20 seconds either predator calls or control sounds were spliced into these playbacks and played for 15 seconds. In total, there were 8 stimuli plus videos without playbacks for experiment 1, and 16 stimuli for experiment 2. The order of stimuli in playlists was edited so that marmosets were not played the same sound sequentially. For the first deployment each group was given the same playbacks in the same order. Further details about the stimuli are given below.

### Experimental stimuli-Predator Calls

Raptors are primate predators (Mcgraw and Berger, 2013) whose calls have been widely used to test anti-predator behaviour and alarm call vocalisations (Fichtel and Kappeler, 2002; Cäsar et al., 2012). Although little is known about pygmy marmoset predation, one report stated pygmy marmosets displayed freezing behaviours and had a higher alarm call rate when they saw large raptors (species not reported) flying overhead (Snowdon and Hodun, 1981). A pilot study was conducted with one pygmy marmoset group which were played the calls of a great black hawk (*Buteogallus urubitinga*), roadside hawk (*Rupornis magnirostris*), ornate hawk eagle (*Spizaetus ornatus*), and a harpy eagle (*Harpia harpyja*). Audio clips were all obtained from the Macaulay Library (see Table SM2 in supplementary materials for the citations of the audio samples used). All four species have been found in the study area (Barker and Papworth, 2024), and three are reported to consume primates (Barnett et al., 2011; de Lyra-Neves et al., 2007; Teixeira et al., 2019), so could elicit anti-predator responses from the pygmy marmoset. Based on the reactions observed in the pilot study group, the ornate hawk eagle and roadside hawk were chosen as the predator conditions for this study. This pilot group was not included in the final experiments.

### Experimental stimuli-Anthropogenic Noise

For the anthropogenic noise conditions there were two motor boat stimuli and two human speech stimuli, all of which were collected in the field. The two audios of human speech were both in English as this is the main language of tourist visitors at the site. One was a conversation between two people and the other was one person speaking. We used these anthropogenic noises as motor boats are the most prevalent man-made noise in the area (Barker, 2022) and human speech is both a good indicator of active human presence and has been used in various playback experiments (Zanette et al., 2023; Crawford et al., 2022).

### Experimental stimuli-Control Audios

The control audio conditions consisted of cicada (*Quesada gigas*) and blue-and-yellow macaws (*Ara ararauna*) both of which are found in the area (Pitman et al., 2003). Both sounds were extracted from YouTube videos and converted to mp3 files (Ambiance - Topic, 2019; Black Crow, 2021).

### Experimental Control- No playback videos

For each group, fourteen videos where a pygmy marmoset was present and there was no playback was selected to serve as a behavioural baseline for the group.

### Video Analysis

All videos from successful playback experiments from the 5 groups were analysed in the software BORIS (version 7.13; Friard and Gamba, 2016), an event logging software for video coding. The pygmy marmoset that was first in frame when the video started became the focal individual for the extraction of behavioural data. When multiple individuals were in frame, we chose to code the adult (for example if there was an adult carrying an infant) or if all were adults then the one which was closer and clearer in the video frame. Within BORIS, an ethogram was created (Supplementary materials Table SM3) which was used to code the videos. The time in frame (TIF) in seconds was calculated for the focal individual for each video. If the focal individual fled, we calculated the time to return in minutes, whether in the same video or when a marmoset triggered the camera trap again. When the marmosets did not trigger the trap again that day (n=2) we limited time to return as the number of minutes until 18:30, as that was when the ABR was set to start the rest period. Two successful playbacks were removed from the final analysis of experiment one as during these playbacks the focal marmoset appeared to be reacting to other aspects of their environment, rather than the playback. Behavioural data for the no playback control videos were extracted in the same way as in the playback videos and included in the dataset for experiment one. Example response to a predator call can be viewed at the following link (https://youtu.be/-j6U4HvcbTo?si=9hy7rQFJPTmBCcUa).

28 videos (22% of the videos analysed) were double blind coded by a primatologist with a background in video coding primate behaviour in BORIS, but without experience working with pygmy marmosets. We found a mean difference of 6.7s per video for eating behaviours (96.3% agreement), 6.9s for vigilance (96.1% agreement), 0.75 vocalisations (99.6% agreement), 0.56s for looking at the camera (99.7% agreement), 0.15s for relaxed behaviours (99.9% agreement), and agreement in 82% of fleeing cases (see Section 2 of the supplementary materials for further details).

### Calculations and Statistical Analysis

The statistical software R (v4.3.2; R Core Team, 2023) was used for all the calculations and statistical analysis described in this section.

### Fleeing Data

A GLMM was run to examine the effect of playback condition on fleeing behaviour. Due to perfect separation (no fleeing was observed in response to the control playbacks), permutation testing (package permutes; Voeten, 2023) with 1,000 iterations was used to calculate empirical p-values for each playback condition.

### Behavioural Differences Between Conditions

As the analysis period was restricted to a two-minute window, the amount of time that an individual could spend on behaviours other than vocalising or fleeing was not independent. To examine potential correlations between behaviours, a PCA (stats; R Core Team, 2023) was conducted on these seven behaviours using the dataset for experiment 1. The behaviours “groom”, “hunt”, “interact” were infrequent and were the only behaviours with negative loadings on the first principle component (PC1). These behaviours were therefore combined into a single dimension, hereafter referred to as “relaxed” behaviours. For continuity and comparability, we also used the combined “relaxed” behaviour when analysing the second experiment. Models were used to assess the relationship between playback conditions and the duration of these five non-independent behaviours (eat, vigilance, looking at the camera trap, looking at the speaker and relaxed). As these five behaviours were not independent, we adjusted the critical alpha for these models to 0.01, using Bonferroni p value correction.

In all models, scaled TIF was included as a covariate, and group was included as a random effect. For each of the five correlated behaviours and vocalisations in both experiments we compared four GLMMs with different distributions; poisson (lme4 package; Bates et al., 2015), negative binomial (lme4 package; Bates et al., 2015) and zero-inflated poisson and negative binomial (glmmTMB package; Brooks et al., 2017). An additional analysis to compare responses to the predation condition in experiment one, to the predator condition when motor boats and human speech were spliced in experiment two was also conducted using a type II Anova. We compared models with the performance package (Lüdecke et al., 2021) and the results of the best fitting model was reported. We excluded models which were overfitting with a conditional R^2^ of 0.9 and above. Where a type II Anova (experiment one) or type III Anova (experiment two) – both run using the car package (Fox and Weisberg, 2019) – indicated there was a difference between conditions in the best fitting model, the GLMM results were examined to assess which conditions differed from one another. Model estimates were extracted using the tidy function (broom package; Robinson et al., 2023).

For the first experiment, a GLMM with a negative-binomial distribution fit was best for “eat” although this did have a singular fit as some groups had identical variation estimates. A negative-binomial distribution was the best fit for vocalisations, “vigilance”, “looking at the camera trap” and “relaxed” behaviours. For the second experiment a negative-binomial distribution was the best fit for vocalisations and the “eat” and “vigilance” behaviour models. For our comparative analysis on the number of vocalisations across the avian predator playbacks in both experiments a negative-binomial distribution was the best fit. The models for some behaviours had a very poor fit, mostly as some behaviours occurred infrequently, so there was a high number of zeros in the response variable. We could not produce satisfactory fits for these models even when using zero-inflated models. These results are therefore not reported here but are included in the supplementary materials for completeness (supplementary materials Tables SM4-8). This was the case for the “looking at the speaker” and “relaxed” behaviours in the first experiment, and the “relaxed”, “looking at the camera trap” and “looking at the speaker” behaviours for the second experiment.

### Ethical Note

The experiment underwent and was approved by the Royal Holloway’s ethical review process. Some of these marmoset groups are in a community, El Chino, and therefore the camera traps could record human conversations and videos of people in the community. We therefore are not sharing all the videos recorded in these experiments, but will share the datasheets with behavioural data extracted from these videos. Permission by the landowners was sought before the camera traps were placed near their homes. The study was approved by the College Animal and Welfare Officer assigned to Royal Holloway, University of London. The research was developed with the permission of the Amazon Research Centre, Peru, meaning no formal research permit is required.

Hearing the predator and human speech audios repeatedly could be distressing for the marmosets. To insure the marmosets did not suffer from prolonged bouts of stress due to the playbacks we only put up the equipment for two days initially, and ensured the marmosets had at least a two-day rest period if the equipment went up again. The 12-minute rest period also ensured the marmosets were not continually exposed to playback trials, and the wide variety of audios on the playlist meant individuals might only hear the predator call or human speech stimuli 4 or 5 times in that period. Our protocol stated that if the marmosets did not return to the active feeding holes during the two days exposure to playbacks we would consider this a significant impact on the study group and re-evaluate the methodology. However, this never occurred and the marmosets always returned to the site after playbacks

## Acknowledgements

A special acknowledgment to Josef Knauseder who designed and created the ABR units used in this project, based on the design of Suraci*et al*. (2017). We would like to thank Dr. Paul Beaver for letting us conduct this study at the site and to the rest of the staff at Amazonia Expeditions who made all the fieldwork possible. Especially Welister Perez Huayta who was our field guide and assisted with the data collection. Thank you to Elisa Fernandez Fueyo who conducted the double-blind coding for this study. We would also like to acknowledge the receipt of media from “The Macaulay Library at the Cornell Lab of Ornithology”.

## Data Availability Statement

Due to the sensitive nature of some of the video files containing people’s private conversations and their image these cannot be given to the public. However, the corresponding author can be contacted in cases of reasonable request for the videos. The data of the behaviours catalogued in each video has been uploaded to Dryad (DOI: 10.5061/dryad.44j0zpcx8) and the analysis script is accessible on GitHub (https://github.com/larissabarker/Monkey-hear-monkey-do-what-an-application-of-ABRsfor-hypothesis-testing.git).

## Supplementary Materials

### Section 1: Referenced tables and figures

**Figure SM1.**
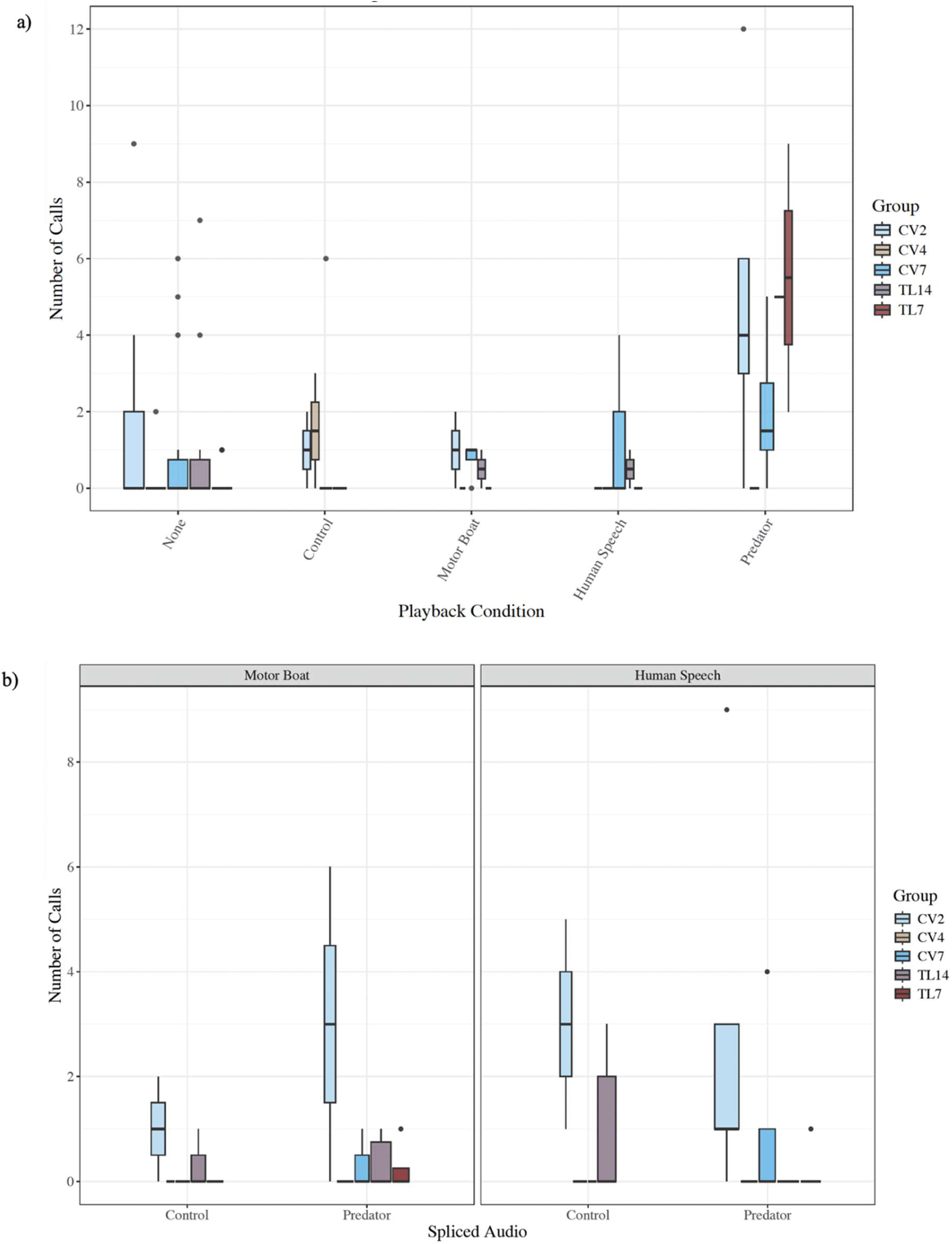
a) the number of vocalisations emitted by the focal individual in experiment one. b) the number of vocalisations emitted by the focal individual in experiment two.

**Figure SM2.**
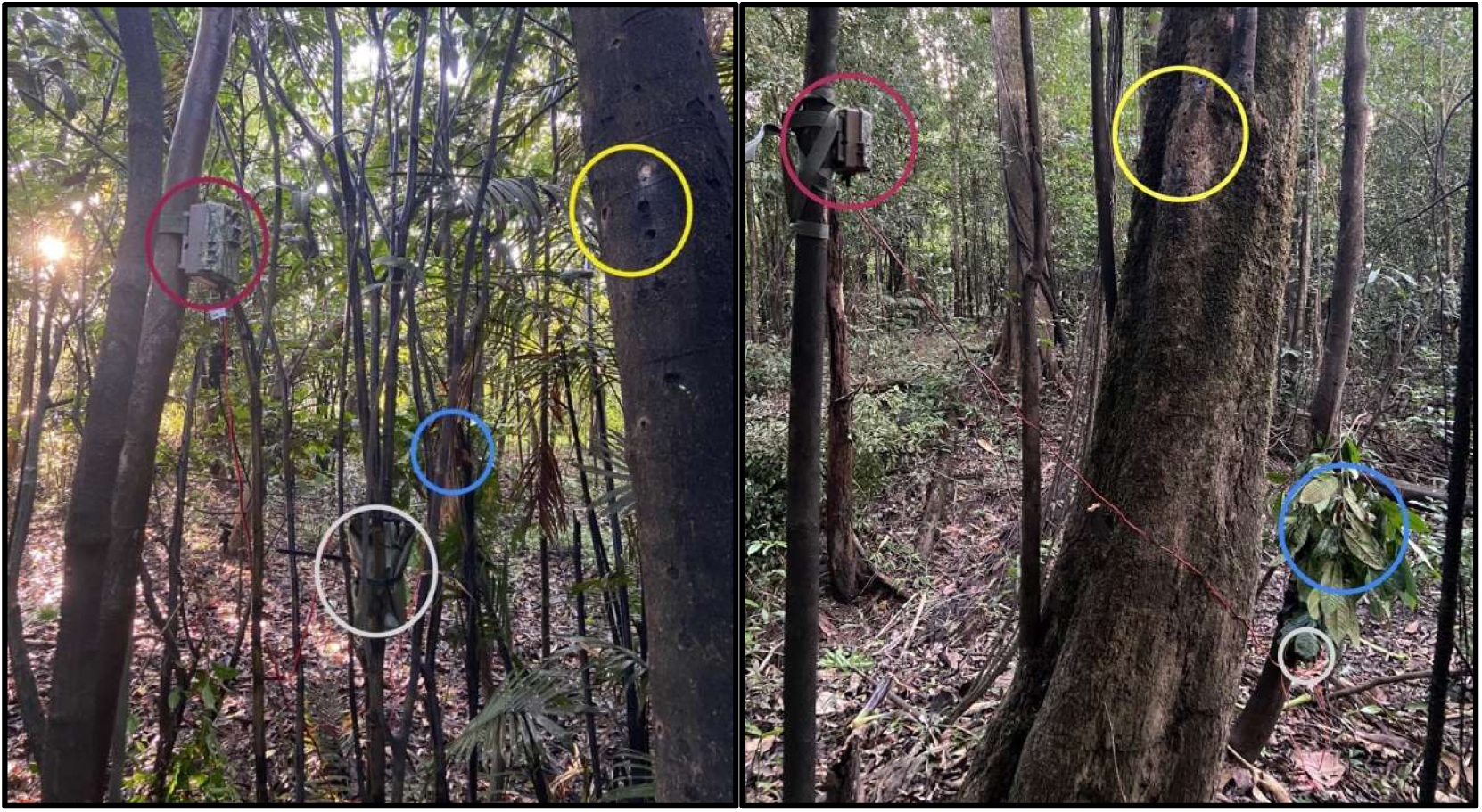
Experimental ABR setup. The maroon circle denotes the position of the camera trap, the grey circle the position of the battery pack, the blue circle the placement of the speaker and the yellow circle highlights the active sap feeding holes. The image on the left was taken at the group TL14 and the right is the group RC2.

**Table SM1.** For both experiments, the time to return after the focal individual fled with the bolded numbers indicating the instances where no pygmy marmoset returned that day.

| Experiment | Playback Condition | Group | Time to Return (min) |
| --- | --- | --- | --- |
| 1 | Motor Boat | TL14 | 2.75 |
|  | None | CV2 | 1.50 |
|  | None | TL7 | 0.30 |
|  | Predator | CV4 | 22.58 |
|  | Predator | CV7 | 16.43 |
|  | Predator | TL7 | 152.73 |
|  | Predator | TL14 | 189.50 |
|  | Predator | TL14 | 32.00 |
|  | Predator | TL14 | 35.53 |
|  | Human Speech | CV2 | 3.35 |
|  | Human Speech | CV4 | 2.60 |
|  | Human Speech | TL7 | **84.80** |
|  | Human Speech | TL7 | 125.57 |
| 2 | Human Speech spliced with Control | CV2 | 0.40 |
|  | Human Speech spliced with Control | CV4 | 12.63 |
|  | Human Speech spliced with Control | CV4 | 32.98 |
|  | Human Speech spliced with Predator | CV2 | 0.67 |
|  | Human Speech spliced with Predator | CV2 | **93.00** |
|  | Human Speech spliced with Predator | CV4 | 4.50 |
|  | Human Speech spliced with Predator | CV4 | 3.30 |
|  | Human Speech spliced with Predator | TL7 | 35.59 |
|  | Human Speech spliced with Predator | TL7 | 1.10 |
|  | Human Speech spliced with Predator | TL14 | 104.62 |
|  | Motor Boat spliced with Predator | CV4 | 1.47 |
|  | Motor Boat spliced with Predator | TL7 | 35.58 |
|  | Motor Boat spliced with Predator | TL14 | 53.48 |

**Table SM2.** Macaulay Library Audio Samples used. The roadside hawk and ornate hawk eagle were chosen for the full experiment due to the marmosets in the pilot having a greater reaction to their calls and the fact they are both known to eat primates or other mammals the size of the marmosets.

| Common name | Scientific name | Samples used |
| --- | --- | --- |
| Great Black Hawk | *Buteogallus urubitinga* | ML13299, ML86013, ML241625 |
| Roadside Hawk | *Rupornis magnirostris* | ML23941, ML80022871, ML165057 |
| Ornate Hawk-Eagle | *Spizaetus ornatus* | ML76017, ML136629, ML75995 |
| Harpy Eagle | *Harpia harpyja* | ML106825751, ML176929541,<br>ML194645771 |

**Table SM3.** The ethogram used in BORIS to code the playback videos.

| Behaviour code | Description | Key | Excluded behaviours |
| --- | --- | --- | --- |
| audio | audio is played/stopped | a |  |
| call | focal individual has made a call, logging calls of all call types | c | eat,groom |
| look | focal individual looks at the camera trap | l | speaker,vigilance,eat,hunt,interact |
| speaker | focal individual looks at speaker | s | look,vigilance,eat,hunt,interact |
| vigilance | focal individual is surveying for threats | v | look,speaker,eat,hunt,interact |
| eat | focal individual is eating sap and or gauging sap holes | e | look,speaker,vigilance,hunt,interact |
| flee | focal individual flees (raid movement out of frame) | f | look,speaker,vigilance,eat,groom,hunt,interac |
| move | focal individual moves position in frame | m | eat,groom,hunt,interact |
| out | focal individual moves out of frame at a normal speed | o | look,speaker,vigilance,eat,groom,hunt |
| groom | focal individual is grooming itself | g | hunt |
| hunt | focal individual is hunting an insect | h | look,speaker,vigilance,eat,groom,interact |
| interact | focal individual is interacting with another pygmy marmoset on frame | i | look,speaker,vigilance,eat,hunt |

**Table SM4.** Experiment 1: Looking at the speaker- poisson distribution without back transformed estimates

| Audio Condition | Estimate | SE | Z | P | N |
| --- | --- | --- | --- | --- | --- |
| None | -20.92 | 156.16 | -0.13 | 0.89 | 70 |
| Control | 19.35 | 156.16 | 0.12 | 0.90 | 12 |
| Motor Boat | 20.24 | 156.16 | 0.13 | 0.90 | 10 |
| Human Speech | 20.83 | 156.16 | 0.13 | 0.89 | 14 |
| Predator | 20.20 | 156.16 | 0.13 | 0.90 | 18 |
| TIF | 0.61 | 0.17 | 3.50 | **0.0005** | 124 |

**Table SM5.** Experiment 1: Relaxed - a negative binomial distribution without back transformed estimates

| Audio Condition | Estimate | SE | Z | P | N |
| --- | --- | --- | --- | --- | --- |
| None | -8.41e<sup>-01</sup> | 8.00e<sup>-01</sup> | -1.05 | 0.29 | 70 |
| Control | -2.60e<sup>+00</sup> | 2.57e<sup>+00</sup> | -1.01 | 0.31 | 12 |
| Motor Boat | -4.08e<sup>-01</sup> | 2.32e<sup>+00</sup> | -0.18 | 0.86 | 10 |
| Human Speech | -6.56e<sup>+01</sup> | 1.79e<sup>+07</sup> | 0.00 | 1.00 | 14 |
| Predator | -3.65e<sup>-01</sup> | 1.79e<sup>+00</sup> | -0.20 | 0.84 | 18 |
| TIF | 1.35e<sup>+00</sup> | 6.60e<sup>-01</sup> | 2.05 | 0.04 | 124 |

**Table SM6.**
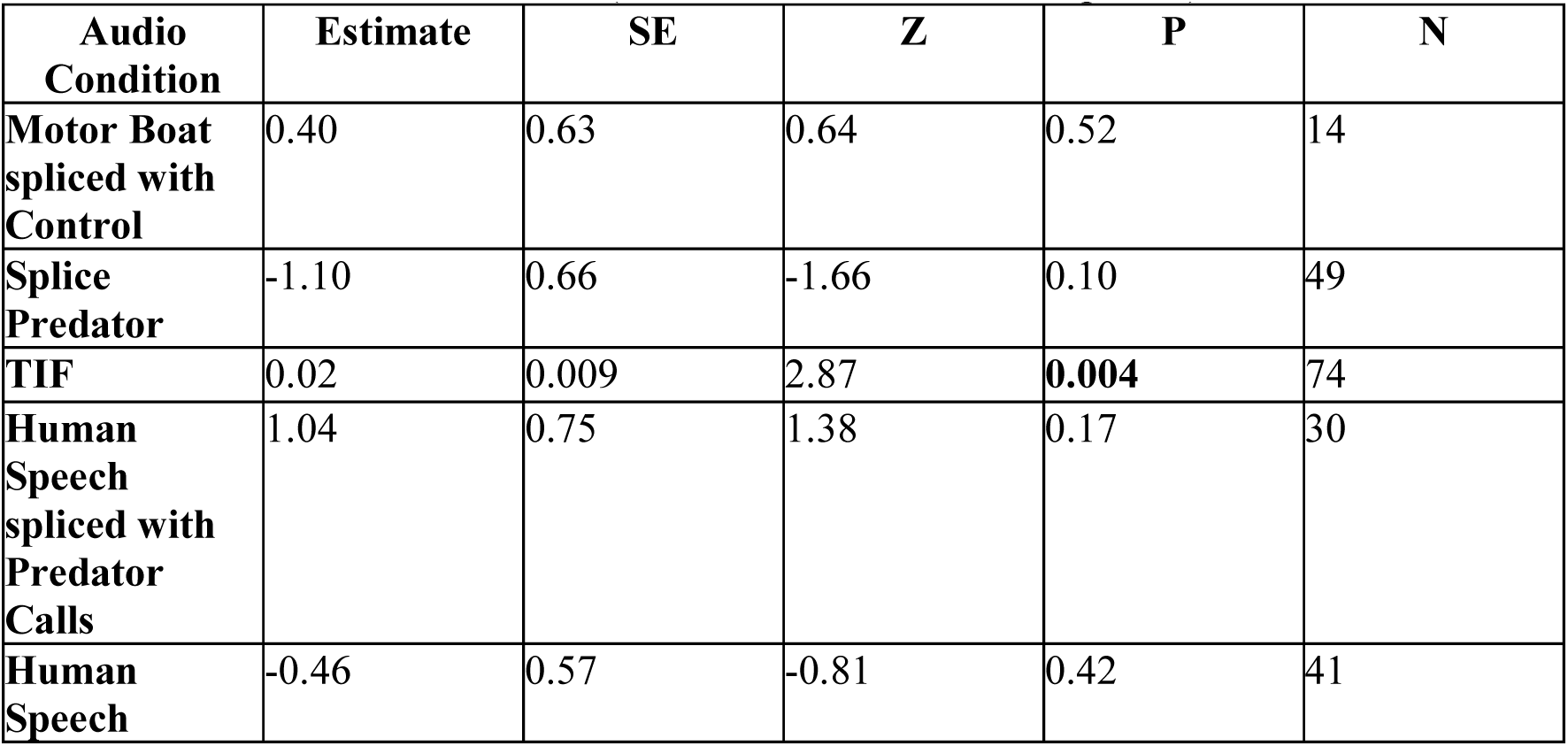
Experiment 2: Looking at the speaker - a zero inflated Poisson distribution without back transformed estimates (conditional model results reported)

**Table SM7.** Experiment 2: Looking at the camera- a negative binomial distribution without back transformed estimates

| **Audio Condition** | **Estimate** | **SE** | **Z** | **P** | **N** |
| --- | --- | --- | --- | --- | --- |
| **Motor Boat spliced with Control** | -0.86 | 0.73 | -1.18 | 0.24 | 14 |
| **Splice Predator** | -0.65 | 0.88 | -0.73 | 0.47 | 49 |
| **TIF** | 0.01 | 0.009 | 1.43 | 0.16 | 74 |
| **Human Speech spliced with Predator Calls** | 0.40 | 1.21 | 0.33 | 0.75 | 30 |
| **Human Speech** | 0.16 | 0.95 | 0.17 | 0.87 | 41 |

**Table SM8.**
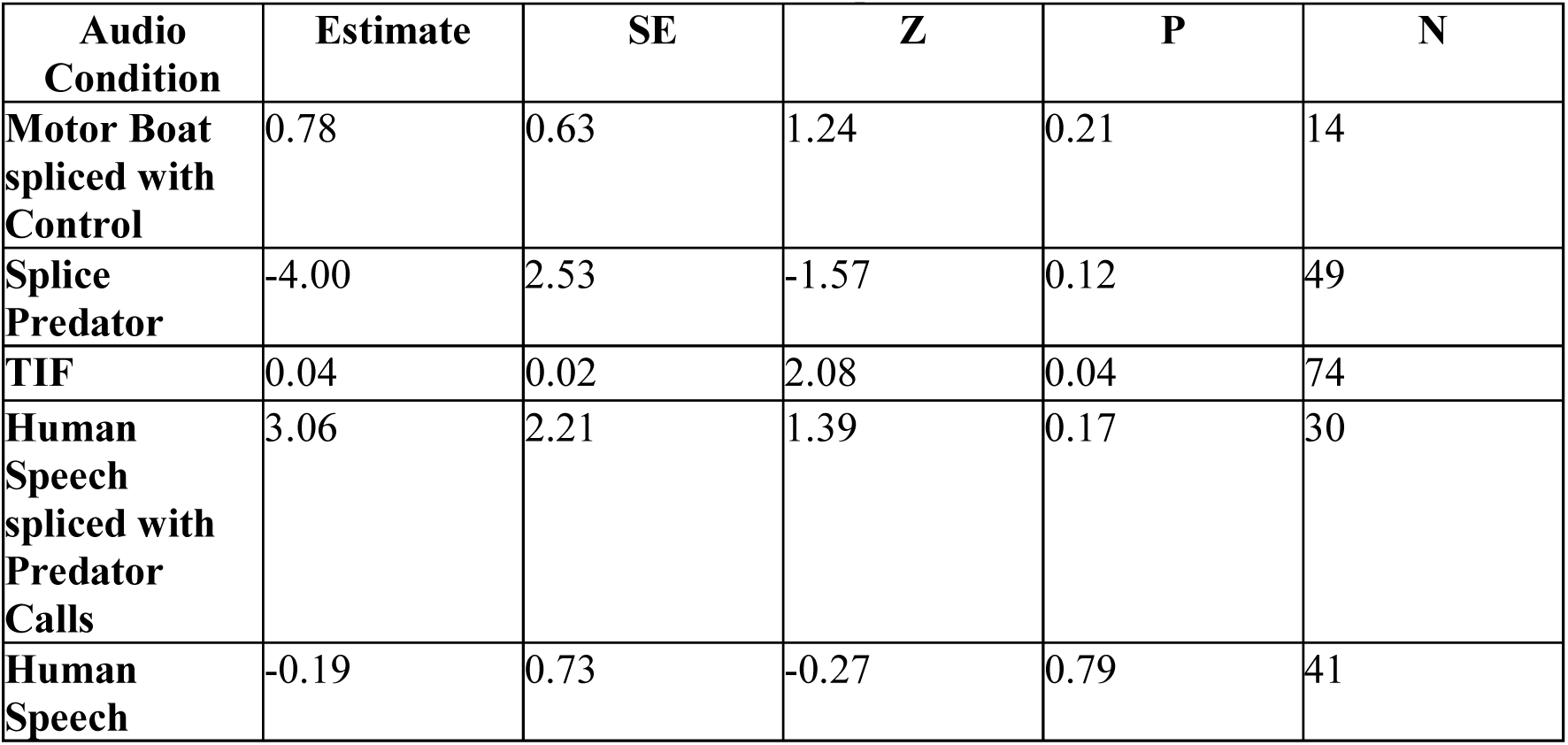
Experiment 2: Relaxed- a zero inflated Poisson distribution without back transformed estimates (conditional model results reported)

### Section 2: Intercoder Reliability

The “looking at speaker” behaviour was not coded for the double-blind coding as the other coder was not familiar with the experimental field setup and the speaker is not always in view in the video. Therefore, we combined “looking at speaker” with “vigilance” for the original coder’s assessment. The sum of each behaviour per video per coder was calculated in R. Then the average difference of these sums for each video was extracted. We calculated the correlation between eating bouts and vigilance bouts and the mean difference in seconds between coders by first testing for normality with Shapiro Wilks tests (stats package; R Core Team, 2023), as neither were normality distributed we ran Spearman’s rank correlation tests (stats package; R Core Team, 2023).

For the percentage agreement we used the average difference per behaviour per video and created the number of seconds the two coders agreed the behaviour was or was not being performed by the focal individual. We then took the average time the coders agreed across all videos for each individual behaviour and extracted the percentage out of the total time of the video (120s).

We had a mean difference of 6.7s per video for “eating” behaviours, 6.9s for “vigilance”, 0.75 calls, 0.56s for “looking at the camera”, 0.15s for “relaxed” behaviours, and agreement in 82% of fleeing cases. Some of these differences are due to the brevity and frequency of these behavioural bouts, where 1 or 2 second differences quickly add up across the 5 or 10 cases of the behaviour per video. We ran Spearmen’s rank correlation tests to show the relationship between the number of bouts of “eating” and “vigilance” behaviours in the video and the mean difference in time recorded by both coders (Figure SM3). These showed a positive and significant correlation for both behaviours. This helps solidify the assertion that the more bouts of a behaviour there is in a video the more likely the coders were to have a greater mean difference in said video. Therefore, we assume the differences between coders is due to small differences in start/stop times culminating over a video rather than differences in behavioural coding.

**Figure SM3.**
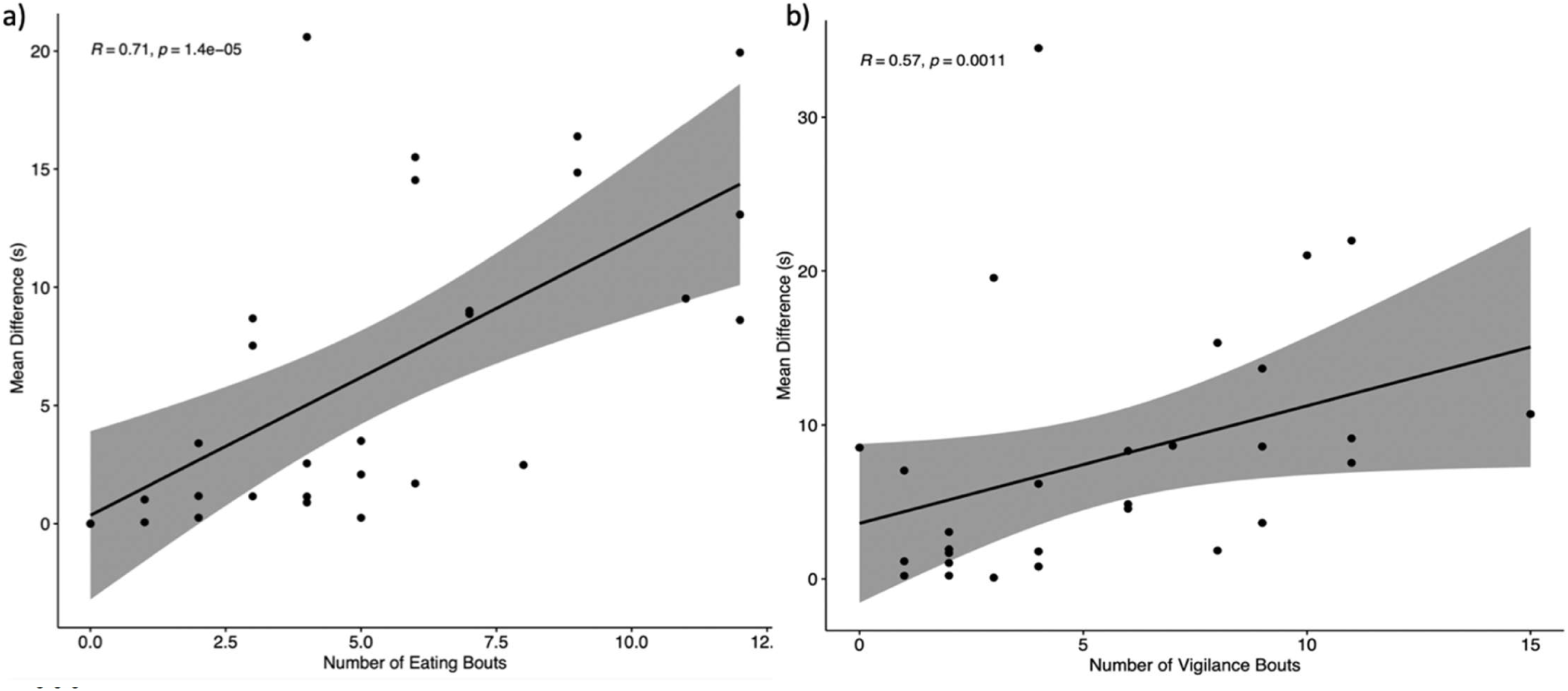
Spearman’s rank correlation between the number of eating bouts (a) and vigilance bouts (b) present in the video and the mean difference per video in seconds.

